# Immunoecology of species with alternative reproductive tactics and strategies

**DOI:** 10.1101/062083

**Authors:** George A. Lozano, Albert F. H. Ros

## Abstract

Alternative reproductive tactics and strategies (ARTS) refer to polymorphic reproductive behaviours in which in addition to the usual two sexes, there are one or more alternative morphs, usually male, that have evolved the ability to circumvent direct intra-sexual competition. Each morph has its own morphological, ecological, developmental, behavioural, life-history, and physiological profile that shifts the balance between reproduction and self-maintenance, one aspect being immunity. Immunoecological work on species with ARTS, which is the topic of this review, is particularly interesting because the alternative morphs make it possible to separate the effects of sex, *per se*, from other factors that in other species are inextricably linked with sex. We first summarize the evolution, development and maintenance of ARTS. We then review immunoecological hypotheses relevant to species with ARTS, dividing them into physiological, life-history, and ecological hypotheses. In context of these hypotheses, we critically review in detail all immunoecological studies we could find on species with ARTS. Several interesting patterns emerge. Oddly, there is a paucity of studies on insects, despite the many benefits that arise from working with insects: larger sample sizes, simple immune systems, and countless forms of alternative reproductive strategies and tactics. Of all the hypotheses considered, the immunocompetence handicap hypothesis has generated the greatest amount of work, but not necessarily the greatest level of understanding. Unfortunately, it is often used as a general guiding principle rather than a source of explicitly articulated predictions. Other hypotheses are usually considered *a posteriori*, but it is perhaps time that they take centre stage. Whereas blanket concepts such as “immunocompetence” and “androgens” might useful to develop a rationale, predictions need to be far more explicitly articulated. Integration so far has been a one-way street, with ecologists delving deeper into physiology, seemingly at the cost of ignoring their organisms’ evolutionary history and ecology. One possible useful framework is to divide ecological and evolutionary factors affecting immunity into those that stimulate the immune system, and those that depress it. Finally, the contributions of genomics to ecology are being increasingly recognized, including in species with ARTS, but we must ensure that evolutionary and ecological hypotheses drive the effort, as there is no grandeur in the strict reductionist view of life.

## 1 INTRODUCTION

At its essence, biological variation is categorical (ACGT), but because of the interacting effects of multiple alleles, genes, and environments, it is usually expressed phenotypically as continuous variation. Nevertheless, discrete, non-continuous variation is surprisingly common. For example, 334 instances of plumage polymorphisms occur in birds, in about 3.5% of all species, distributed across 53 of the 143 families, and 14 of the 23 orders (Galeotti et al. 2003). In insects, wing polymorphisms are common (Roff and Fairbairn 1991; Zera and Denno 1997), and polymorphisms producing distinct castes (O’Donnell 1996) are one of the best-known characteristics of eusocial insects. Of all the types of discontinuous variation, morphological variation is perhaps the most readily evident, but it is usually associated with discontinuous variation in life history and behaviour, all of which is, of course, regulated physiologically. For example, insects’ wing polymorphisms (Roff and Fairbairn 1991) are associated with dispersal polymorphisms (Zera and Denno 1997) and mediated by juvenile hormone (Dingle and Winchell 1997). A wide assortment of other such polymorphisms exist, for example, in chemosensory recognition (López et al. 2009), foraging (Ehlinger and Wilson 1988), fighting (Crespi 1986) and in reproduction (Oliveira et al. 2008b). In this review, we examine immunity in species with a specific type of polymorphic reproductive behaviours.

Alternative reproductive tactics and strategies (ARTS) refer to polymorphic reproductive behaviours in which in addition to the usual two sexes, there are one or more alternative morphs, usually male, that have evolved the ability to circumvent direct intra-sexual competition. Unfortunately, the ARTS literature has become a definitional quandary because as it developed, it relied on colloquial terms that were given additional and special meanings. Semantically, “strategies” are simply long-term “tactics”, and “tactics” are nothing more than short-term “strategies”, big picture versus detailed plans, relatively immutable versus highly variable. In the evolutionary ecology literature, the term “strategy” was initially used to refer to variation resulting from genetic effects, and the term “tactic” to variation resulting from environmental factors (Alcock 1979; Maynard-Smith 1982). This nomenclature carries with it the implicit assumption that genetic effects are permanent and environmental effects are ephemeral. Although this way of thinking is outdated and unnecessary (Brockmann 2001; Oliveira et al. 2008a), the association still exists in the literature and in people’s minds.

Similarly, some authors use the term “polymorphic reproductive behaviours” synonymously with “alternative reproductive behaviours” (reviewed by Oliveira et al. 2008b). However, the term “polymorphic” is more general, literally only implying the existence of more than one form. In contrast, the adjective “alternative” implies that all options are not equal, and that some of them are not the conventional or mainstream, as in “alternative lifestyle” or “alternative music”. In this sense, the term “alternative reproductive behaviour” actually refers to reproductive behaviour that is absent in phylogenetically or ecologically related species, which only have the “normal” behaviour. For example, in the halictid bee *Lasioglossum erythrurumoccur* (Houston 1970; Kukuk and Schwarz 1988) and the andrenid bee *Perdita portalis* (Danforth 1991; Danforth and Neff 1992), “normal” males leave the nest in search of mates, as in related species, but in both species there are also “alternative” males. These are large and flightless males that do not leave the nest but instead fight for access to females within the nest. Before ARTS were conceptually unified, they were recognized and described in many species. Therefore, a variety of names have been used to refer to the “normal” and “alternative” morphs, such as independent and satellites, fighters and scramblers, fighters and sneakers, bourgeois and kleptogamic, phallic and aphallic, bourgeois and parasitic, M+ and M-, producers and scroungers, etc., but in many respects, these pairs of terms are functionally equivalent.

Several authors have tried to clarify this nomenclature (Caro and Bateson 1986; Brockmann 2001; Oliveira et al. 2008a), evidently to no avail, probably because of the pervasiveness of the colloquial sense of the co-opted words, the need for continuity, and, as always, intellectual inertia. Here, we will explicitly define the terms we use and the scope of this review. First, we use the term “alternative reproductive behaviours” to mean a specific type of polymorphic reproductive behaviours in which there is a conventional, “normal” reproductive behaviour, and one or more alternatives. The “normal” behaviour is found in related species in which the “alternative” does not exist. The alternative has been referred to as the “disfavoured role” (Uglem et al. 2001), but it does not have to be so. The alternative might actually be the main or primary reproductive behaviour (Almada and Robalo 2008). For example, in the fallfish minnow *(Semotilus corporalis)*, less than 10% of mature males are territorial and the rest as sneakers (Ross 1983). Second, we will not place any limitations on the underlying mechanisms, genetic or environmental, but we shall distinguish whether the reproductive behaviours and morphs are fixed for the individual’s entire lifetime, or plastic, when individuals change from one to another (Rivas-Torres et al. 2019). If plastic, we will make the distinction of whether the change occurs only once, or the change(s) is/are reversible (Moore 1991; Moore et al. 1998; Oliveira et al. 2008a). In most cases, we will be addressing cases in which these alternative reproductive behaviours are relatively long-term, at least sufficiently so to affect immunity, but to avoid the dilemma of when exactly a tactic becomes a strategy and vice-versa, we shall use the more inclusive term “alternative reproductive tactics and strategies” (ARTS).

Species with ARTS have at least one additional life-history pathway that is not present in other species. These alternative reproductive pathways require a new set of central and peripheral adaptations that in other species are strictly associated with each sex. Hence, species with ARTS make it possible to separate the effects of sex *per se* from the physiological, morphological, or behavioural differences between the morphs, providing additional combinations of reproductive traits within each sex (Noble et al. 2013; Marson et al. 2019).

In this paper, we first briefly address the evolution, maintenance, and development of ARTS, relying mostly on several excellent reviews (Brockmann 2001; Oliveira et al. 2008a; Taborsky et al. 2008), and focusing primarily on areas specifically relevant to immunoecology. Second, we examine several ultimate and ecologically-based proximate hypotheses that predict differences in immune function between the sexes, and by extension, among morphs in species with ARTS. Third, we review in detail immunoecological work on species with ARTS, specifically relating it to the aforementioned hypotheses. Finally, we offer prospects for future work, both for researchers already working with species with ARTS, researchers working on immunoecology, and of course, to guide our own future endeavours.

## 2 EVOLUTION, MAINTENANCE, AND DEVELOPMENT

We include a brief summary of the evolution, development and maintenance of ARTS. This section should not be considered a comprehensive review; several are available (Gross 1991; Tabosrky 2001; Oliveira et al. 2008b; Taborsky and Brockmann 2010). The purpose here is merely to establish some common ground to facilitate the ensuing discussion.

### 2.1 Origin and maintenance of ARTS

The stage is set up for the evolution or ARTS when a high population density coupled with intense intrasexual selection allow only the largest and strongest males to obtain most of the copulations (Kokko and Rankin 2006). Interactions among males might include territorial defence, direct confrontations, and extended and/or elaborate sexual displays. In such situations, males are selected to divert resources towards growth and, given that they are unable to reproduce while they are small, perhaps even to delay sexual maturity. On the other hand, this scenario also favours adaptations that allow males to reproduce while avoiding this intense competition. Contrary to popular belief, males seldom “want a challenge” when it comes to mating. Males are usually quite willing to mate opportunistically, without having to search for females, compete with other males, establish and defend territories, court females, or pay any of the costs associated with reproduction.

ARTS are long-term, sometimes permanent, distinct phenotypes specializing in opportunistic low-cost mating. Because these specializations are relatively long-term to permanent, ARTS have evolved along with a wide range of similarly long-term to permanent behavioural, morphological, and physiological adaptations.

We usually observe the result of evolution but seldom do we have the privilege of witnessing evolution in action. When ARTS first begin to evolve in a population, avoiding intra-sexual competition is only the first step. Before the alternative male phenotype is established, females are under strong selection to avoid mating with small, subordinate males. Hence, for “alternative” males to be successful, they must not only avoid direct competition with “normal” males, but also bypass female choice. In many cases, they do both via sneak copulations. However, once the alternative morph is established, females would no longer be selected to avoid these males; in fact, the loci for male morph and female preference for that morph would be linked (Wellenreuther et al. 2014). Thereafter, female preference for the alternative morph might fluctuate, depending on the morph proportions and their relative reproductive success. A final requirement is disruptive selection, whereby intermediate males are disfavoured, both because of female avoidance (Cummings and Ramsey 2015) and because of their inability to compete with either of the two other types of males (Fig. 1).

**FIGURE 1.**
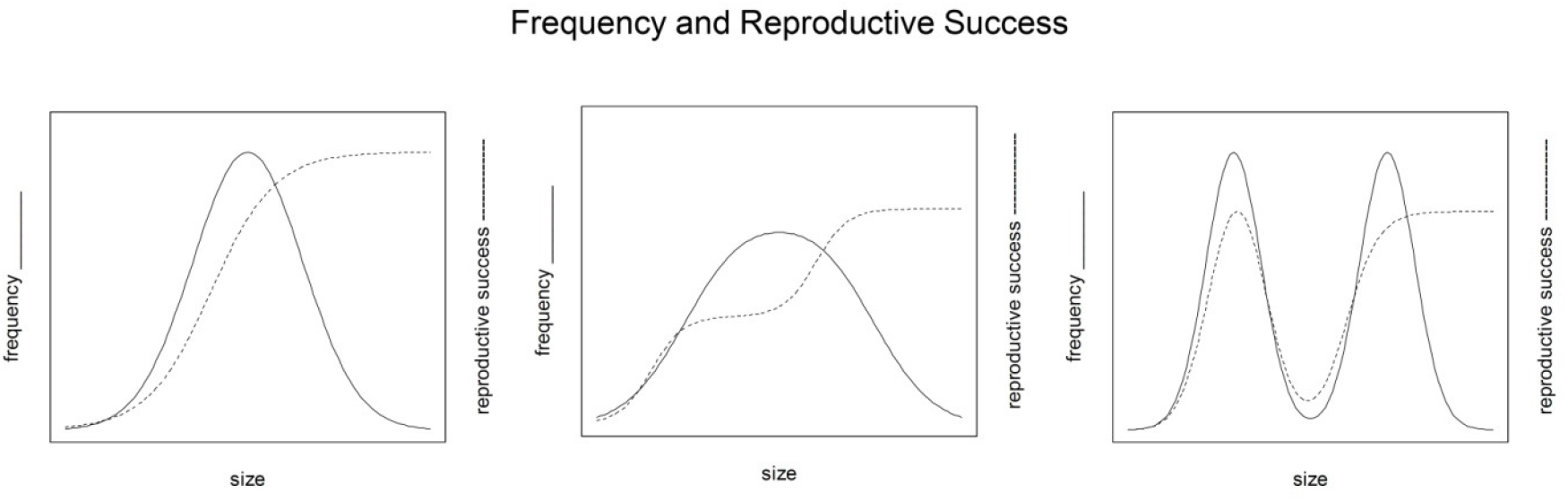
Frequency and reproductive success of “normal” and “alternative” males during the evolution and establishment of ARTS in a population. “Size” is used in the x-axis, but it could also be any other aspect of behaviour or morphology that is important in mating. At the start, on the left, there is only one morph with a highly skewed distribution in reproductive success. In the third graph, there are two distinct groups. The final relative frequencies of the 2 morphs might vary depending on the equilibrium that is eventually reached.

In species with ARTS, each morph takes a different approach to obtaining copulations, so antagonistic selection (Holland and Rice 1998) does not occur just between two sexes, but also separately between each morphs. In due time, through condition-dependent and pleiotropic effects, genetic correlations develop between traits, whether or not these traits are functionally related (Lawson et al. 2011; Kupper et al. 2016). These sets of traits and their interrelationships often have a long evolutionary history, and striking examples occur in sexual dimorphic sets of behavioural and morphological traits (Lande 1980; Wellenreuther et al. 2014; Barron et al. 2015). Furthermore, intralocus sexual conflict, whereby different optima for each sex exist within a locus, is extended to include intra-locus conflict for different morphs within one sex (Morris et al. 2013; Bielak et al. 2014).

On an evolutionary time scale, ARTS are relatively ephemeral; they do not become permanent features of relatively large clades. For example, a phylogenetic analysis of blennies, wrasses, and salmonids indicates that ARTS evolved independently in several lineages, but they also readily disappeared (Almada and Robalo 2008). In the side-blotched lizard, *(Uta stansburiana)*, described in more detail below, there are 3 male morphs. The proportion of these 3 morphs varies geographically, and all morphs are not present in all populations. Territorial morphs can continue to function without sneakers, but sneakers cannot function in the absence of territorials. So, whenever one of the morphs is absent, it is always the sneaker morph. Allelic frequencies immediately shift when sneakers are lost in a population, causing rapid morphological divergence, and leading to distinct populations, or even sub-species (Corl et al. 2010). The disappearance of polymorphisms is one possible mechanism of speciation (West-Eberhard 1986).

Whether permanently or not, the maintenance of ARTS can be considered in the same context as, or as an extension of, sex allocation theory (Darwin 1871; Fisher 1930; Charnov 1982). Essentially, a negative frequency-dependent system is set-up whereby the alternatives accrue differential benefits that depend on each other’s frequency in the population and/or their respective fitness. Costs and benefits of each morph depend on population dynamics, such as the changes in the density and proportion of morphs in a population. Genetic variation between and within morphs might code for developmental pathways leading to ARTS, and for responses to environmental variation that specifically take into account morph proportions. In both cases, ARTS might be selected against when the balance between costs and benefits gets tipped in favour of a single phenotype. Therefore, to understand the maintenance of ARTS, it is important to study not only the genetics, but also the costs and benefits of the associated traits (Oliveira et al. 2008a)

### 2.2 Development of morphs

Depending on the species, morphs develop via a variety of hormonal mechanisms (Rhen and Crews 2002). ARTS can be fixed or flexible; if flexible, they can be reversible or not. These options evolve depending on the mating opportunities, and the costs and benefits involved in switching back and forth. The benefit of fixed morphs is that they can be more specialized, behaviourally, morphologically, and physiologically, and that specialization can begin to take shape as soon as one developmental path is chosen over the other. The drawback of a fixed morph is the loss of opportunities, which may be more important when mating opportunities vary widely spatially or temporally. Hence, reversible choices are more likely to occur in adults and take place relatively quickly, in a matter of days, whereas and non-reversible choices occur more often at an earlier developmental stage and their development take relatively longer (Emlen 2008).

Whether they are reversible or not, when morph changes are plastic, they develop via threshold mechanisms. These mechanisms entail monitoring the environment, biotic and abiotic factors, particularly the density and frequency of conspecific morphs, and comparing that information against some internal threshold to choose whether to start changing. All aspects of the process are regulated by hormones. The rate of hormone synthesis, secretion, and degradation, and the expression, production, and binding affinities of receptors all vary and are under selection (Emlen 2008). In insects, these changes are usually regulated by juvenile hormone (Brockmann 2008), and in vertebrates, by sex steroids. One-time changes are more both onerous and carry more longterm consequences than reversible changes, hence they are under stronger selection. In the few cases in which morph determination is strictly genetic, peripheral adaptations can begin to develop as early as possible. In fact, genes associated with a given morph are expected to be linked.

In vertebrates, hormonal control of sexual development traits is usually separated into organizational effects, which occur early in development, and activation effects, which occur during adulthood (Phoenix et al. 1959). For example, the sexual differentiation of the brain and genitalia, which begin before birth or hatch, are organizational events, whereas changes in behaviour and morphology that occur every breeding season are activational events. In addition, there is puberty, a period of both re-organization and activation (Romeo 2003). This paradigm has been extended to the development of morphs in species with ARTS (Moore 1991), the suggestion being that regulation is relatively more important in species with fixed morphs, and activation relatively more important in species with plastic morphs. However, a survey of the literature fails to find support for this hypothesis (Oliveira et al. 2008a). A thorough analysis indicates that the situation is complex: depending on the species, several biochemically related androgens are involved in the development of ARTS, in different proportions and with different effects. One generalization that does arise is that androgens are more important in the development of morphological ARTS than in the development of strictly behavioural ARTS (Oliveira et al. 2008a).

Finally, in the standard vertebrate, sex hormones released by the gonads not only regulate reproduction but also morphology and behaviour. In species with ARTS, sex hormones regulate gametogenesis in both morphs, but their effects on morphology and behaviour differ between morphs. Therefore, it is important not only to examine the production, release, and distribution of various hormones, but also their effects on target tissues.

## 3 IMMUNOECOLOGICAL HYPOTHESES RELEVANT TO SPECIES WITH ARTS

The immune system protects animals against parasites. Parasites are defined here broadly and functionally to include organisms that live in or on a heterospecific animal, the host, obtain nutrients primarily from the host, and have the potential to decrease its fitness (Lozano 1998). Pathogenicity can vary, and parasites often coexist with their hosts without causing any measurable deleterious effects most of the time, but can suddenly overwhelm immunocompromised hosts (Walzer and Genta 2020; Schmid-Hempel 2021). Resistance to parasites is often measured by the “prevalence”, the number or proportion of hosts in a population infected by a given parasite, and “intensity”, the mean number of parasites in individual hosts. Like all other physiological processes, immune function carries an energetic cost. This cost has been confirmed by direct energetic measures (e.g., Barr et al. 1922; Demas et al. 1997; Muehlenbein et al. 2010) and by examining trade-offs between immune function and other energy-demanding processes (e.g., Allander and Bennett 1995; Xu and Wang 2010; Marais et al. 2011). This concept of trade-offs is pivotal to the study of immunity in an evolutionary ecology context, and the basis for several ultimate and proximate hypotheses detailed below. This section presents these hypotheses and their predictions, in the context of species with ARTS. These hypotheses do overlap and hence, are not mutually exclusive, but for the purposes of this discussion, they are divided into physiological, life history, and ecological hypotheses.

### PHYSIOLOGICAL

#### 3.1 Immunocompetence handicap hypothesis (ICHH)

The ICHH posits that honesty in vertebrate sexual signals is maintained because testosterone, the hormone that causes the development of sexual signals in males, also has the effect of depressing immune function (Folstad and Karter 1992). Hence, a trade-off between display and immune protection is expected, and males who can afford to depress their immunity are better able develop more prominent sexual displays. The honesty of these sexual signals (sensu Zahavi 1975) is guaranteed by their immunological cost. Although only testosterone and only males were mentioned in the original hypothesis, in its general form, the hypothesis also applies to several other androgens, and to females to the extent that they are exposed to androgens.

Tests of the ICHH have been correlational and experimental. Correlationally, males should be immunosuppressed relative to females, more so at times of testosterone peaks during their breeding cycles. Experimentally, exposure to testosterone should enhance sexual traits in males and decrease immune function. Conversely, an experimental decrease in testosterone should depress the development of sexual traits and improve immune function. Of course, the timing of any such manipulations is important. Depending on the species, one would have to consider the timing of the development of seasonal sexual traits, and the timing of the breeding season itself. Nevertheless, in both instances of experimental exposure to testosterone, organisms would be moved away from their individual optima, and regardless of the effects on immunity or sexual signals, this imposed change should have an overall negative effect on fitness. In general, studies fail to find support for the key prediction that testosterone depresses immunity (reviewed by Roberts et al. 2004; Bejarano and Jahn 2018).

In the ICHH, the exact nature of the link between immunity and sexual traits depends on the specific effects of testosterone. The hypothesis was based on the physiological effects of testosterone, but did not incorporate the mechanisms behind these effects. One possibility is that testosterone explicitly depresses specific aspects of immunity, perhaps then secondarily liberating resources for other functions, including sexual displays. Another not necessarily mutually exclusive mechanism is that testosterone forces the development of sexual displays, in the process using up resources and energy that would otherwise be allocated towards other biological functions, including immunity (Wedekind and Folstad 1994).

Using testosterone-exposed females, the two alternatives could be tested. One option predicts that immune function should decrease upon experimental exposure to testosterone; the other option predicts that immune function would decrease only if testosterone-exposed females develop male-like sexual ornaments or engage in costly male-like behaviours. Of course, exposing females to unnatural levels of male hormones might have unknown and undetected consequences that would make interpretation difficult. In contrast, using species with ARTS it is possible to compare immune function among different types of males, each with its distinct hormonal profile (Ketterson and Nolan 1999).

The ICHH’s predictions differ depending on the presence of testosterone, its distribution over the animal’s body, and the sensitivity to it of various body tissues. For example, in female mimicking morphs, spermatogenesis still depends on testosterone, but circulating levels might differ from those of “normal” males, and various tissues and organs might be differentially sensitive to it. Hence, in some species with ARTS, it is possible to separate the direct effects of testosterone *per se* on immunity, from the indirect effects caused by a reallocation of resources towards sexual displays.

Similarly, the ICHH generates different predictions in species with ARTS in which the morphs are reversible versus species in which the morphs are permanent. In reversible morphs, one would expect fluctuations in androgens and immunity that reflect a more cautious approach, only a partial commitment to a given morph. In contrast, individuals have more at stake when the change is permanent, and as a terminal gambit, final stage males might expose themselves to high levels of androgens, and be more willing to suppress their immunity to allocate more resources towards reproduction.

A related and supposedly competing hypothesis has also been proposed: the “immunodistribution hypothesis” (Braude et al. 1999). One way immunocompetence is measured is by counting various immune cell types present in the blood. The immunodistribution hypothesis argues that these measures might give spurious results because testosterone might not actually decrease the numbers of any specific cell type, but rather merely redistribute these cells differentially throughout the body, and this redistribution might then be mistakenly interpreted as immune-suppression (Braude et al. 1999). Unfortunately, the immunodistribution hypothesis does not specify what happens to these cells after they are no longer circulating, and nor the extent to which different cell types are necessary in the blood stream as opposed to other tissues. Hence, this hypothesis is more of a critique of a specific method of assessing immunocompetence, whether or not testosterone is involved, rather than an alternative to the ICHH.

The opposite mechanism, that immune activation depresses testosterone, has also been proposed (Boonekamp et al. 2008), and it also predicts that only healthiest males should develop the most elaborate secondary sexual traits (Table 1). This mechanistic hypothesis differs from the ICHH only in the directionality of the effect; the ICHH says that testosterone affects immunity, and this one says that immune activation lowers testosterone. However, this distinction yields different predictions and different implications about the meaning of sexual signals to females.

**TABLE 1.** Contrasting predictions about the relationship between sexual signals and immunity, depending on whether testosterone lowers immune function (Folstad and Karter 1992), or the reverse mechanism, that immune activation lowers testosterone (Boonekamp et al. 2008).

| *Experimental Treatment* | *T lowers immunity* | *Immunity lowers T* |
| --- | --- | --- |
| Increase in T | (1) Decreased immunity.<br>2) More elaborate sexual traits. | (1) No direct effect on immunity.<br>(2) More elaborate sexual traits.<br>(3) If resources are lacking, a decrease in all other energy requiring functions, not just immunity. |
| Immune challenge | (1) No effect on T.<br>(2) Decreased sexual traits-due to energetic requirements. | (1) Decrease in T.<br>(2) Decreased sexual traits-due to energetic requirements and lower T. |
| Information gleaned from sexual traits | Willingness to increase T and risk disease at the time sexual traits developed. | Low exposure to parasites at the time sexual traits developed. |

#### 3.2 “Sperm protection” hypothesis

Another testosterone-based hypothesis, the sperm protection hypothesis, states that testosterone suppresses immunity because spermatocytes are haploid and recognized as “foreign” by the immune system (Hillgarth et al. 1997). Spermatocytes are located within one of several immunoprivileged areas in the body (Streilein 1996), where they are physically isolated from direct contact with blood. Nevertheless, some lymphocytes might squeeze through the testes-blood barrier, and spermatocyte antigens might escape and enter circulation. The suggestion is that to prevent the attack of sperm cells, testosterone, in addition to inducing spermatogenesis, also causes a local suppression of immunity. Furthermore, the hypothesis argues, testosterone causes immunosuppression elsewhere in the body when it either spills over or when it is necessary for purposes other than spermatogenesis.

Few explicit test of this hypothesis exist, but a meta-analysis of 32 studies in which corticosteroids were used to treat infertility in men showed that corticosteroid reduced plasma and seminal anti-sperm antibodies, increased sperm count and motility, and ultimately increased pregnancy rates (Skau and Folstad 2005). Other than the predicted effect of testosterone specifically on areas near the testes, the predictions of this hypothesis do not differ from those of the ICHH. Both hypotheses argue that testosterone is necessary for the development of secondary sexual traits, and is also immunosuppressive, so a trade-off occurs and hence, sexual signals are guaranteed to be honest. The sperm protection hypothesis, however, does provide a functional reason as to why testosterone lowers immunity.

Although this hypothesis has not received much detailed attention, it might be particularly applicable to species with ARTS. One general prediction is that animals with large and prolific testes would be relatively immuno-suppressed. In several species with ARTS, one male morph (dominant, large, pugnacious, and/or territory holders) displays the typical somatic effects of testosterone, and the other morph (small, sneakers, non-territorial, and/or female mimics) sometimes has disproportionably large testes. Hence, correcting for the differences in somatic and gonadal sensitivity of testosterone, and the relative effectiveness of the respective blood-testes barriers, it might be possible to test whether immune function decreases because of a spill-over of testosterone, or because of a reallocations of resources towards growth.

### LIFE HISTORY

#### 3.3 Bateman’s Principle

A limitation of the ICHH and its derivatives is that they apply only to species in which sexual ornaments depend on testosterone, namely vertebrates. However, sex differences in immunocompetence should also be expected simply based on life history (Rolff 2002). Variance in reproductive success is usually greater for males, and generally males maximize their reproductive success by mating with more females, whereas females do so by increasing their fecundity by having longer reproductive lifespans (Bateman 1948). Hence, life history predicts that investment in immune function ought to be greater and less variable for females than for males. More generally, extending the logic to species with ARTS, the expectations are that the sex, gender, or morph with greater lifetime variance in reproductive success should have a more risky lifetime strategy and hence investment on immune function should be lower and more variable.

A strict life-history hypothesis would still be viable in the absence of a testosterone-mediated system. The rationale does not depend on the mechanism regulating the trade-off, so it is applicable to non-vertebrate taxa, and even non-animal taxa. Non-animal taxa might not have an immune system *per se*, but still invest on pathogen defence. Another important difference is that from a life history perspective, sex differences in immunity are expected before the onset of reproduction and would work and even in the absence of costly sexual ornaments in males. The moment sexes or morphs are determined, investment in immune function would start to follow different paths.

### ECOLOGICAL

#### 3.4 Energy and resource constraints

The basic premise of life-history theory is the fact that resources are limited and organisms must face choices in the allocation of resources towards various functions (Williams 1966; Pianka and Parker 1975; Stearns 1992; Roff 2002). All physiological processes carry an energetic cost. Hence, everything else being equal, additional energetic requirements, such as sexual displays, reproduction, thermoregulation, territorial defence, or parental effort, might reduce investment in immunity. Of course, everything else is seldom equal, but nonetheless, a wide variety of trade-offs involving immune functions have been examined. For example, eliciting an immune response increases energetic costs (Marais et al. 2011); fasting suppresses immune responses (Xu and Wang 2010); thermal stress depresses immunity (Dabbert et al. 1997), and higher reproductive effort decreases long-term immune function (Ardia et al. 2003). Countless other examples exist.

However, finding the expected negative relationship between two resource- or energy-requiring functions is not particularly enlightening. Given limited resources, such dual trade-offs are a physical necessity. A more interesting scenario emerges when more than 2 players or energy-requiring functions are examined concurrently. For example, like any other animal, a reproducing female is expected to avoid parasites (Hart 1990). However, when a third party is involved, her offspring, and given that antibodies are transferred to offspring (Hasselquist and Nilsson 2009), she could also benefit by mithridatically^1^ exposing herself to parasites, developing immunity, and transferring that immunity to her offspring (Lozano and Ydenberg 2002). Similarly, when morphs are involved, not just the 2 sexes, the behavioural, ecological, and evolutionary options become more interesting. Some male morphs might spend energy providing paternal care, developing sexual signals that also attract predators, or constructing structures for mate attraction and brood care (Andersson 1994). In species with ARTS, “alternative” males pay only some of the costs that “normal” males pay, so these species offer more appropriate systems in which to further test these trade-offs.

#### 3.5 Exposure Risk

Behavioural defences against parasites come in many forms, including, for example, the physical removal of parasites (Murray 1987; Grutter 1996), prophylactic or therapeutic self-medication (Lozano 1998; Villalba et al. 2010), and the avoidance of certain habitats (Piersma 1997), food items (Keymer et al. 1983; Fleurance et al. 2007), or individuals (Thomas et al. 1996; Ramnath 2009).

Similarly, immune defences can also vary depending on the habitat (Lindström et al. 2004; Ardia 2005), and the expected level of parasite exposure (Joop and Rolff 2004). Seasonal changes in immune function (Nelson 2004) suggest that animals can pre-emptively adjust their immune systems in anticipation to parasite exposure (O’Neal 2013). Hence, differences between the sexes, or morphs, might exist even in the absence of any of the effects proposed by the aforementioned hypotheses. Given differences in behaviour, whether they are differences in foraging, migration, mate searching, mating, parental care, etc., morphs might be differentially exposed to parasites and would be expected to invest on their immunity accordingly.

## 4 IMMUNOECOLOGY IN SPECIES WITH ARTS

### 4.1 Fish

Among vertebrates, ARTS are particularly prominent in fish because of several reasons. First, external fertilization is the norm in fish, which makes it more difficult for males to control access to females, and for females to enforce their partner choice. Both these factors facilitate the evolution of sneakers. Second, fish often have indeterminate growth, so in some cases adult males have to compete with males several times their own size, which favours the evolution of reproductive tactics and strategies that do not rely on direct competition. Third, flexible and variable sex-determination mechanisms allow for the evolution of age-, size- and condition-dependent ARTS, with some fish being able to change their sex and male morph back and forth. Finally, when parental care occurs in fish, it is more likely to be paternal, not maternal or bi-parental, a condition that favours the evolution of parasitic males that exploit other males’ paternal care. Hence, the variety and complexity of ARTS among fish is unlike that of other vertebrate classes (Taborsky 1994; Fleming 1996; Tabosrky 2001; Garant et al. 2003; Taborsky 2008; Wood et al. 2022), but surprisingly, their immunoecology has only been examined in a handful of species.

**The Arctic char** *(Salvelinus alpinus)* is a circumpolar Salmonid that breeds in fresh water. Their life histories are quite variable. They can be either landlocked or anadromous; their final adult size ranges from 13 to 75 cm, and the age of maturity from 3.5 to 10 years (Vøllestad and L’Abée-Lund 1994). During the breeding season, both sexes display carotenoid-dependent red coloration in their abdomens (Scalia et al. 1989; Martinkappi et al. 2009). Depending on their relative size in their respective populations, males use one of two mating tactics: guarding or sneaking. Females are aggressive towards small sneakers, which could be interpreted as a preference for larger males, or a defence against nest predation. In addition, some populations have a dwarf male morph that is exclusively a sneaker (Sigurjónsdóttir and Gunnarsson 1989). Compared to guarding males, sneakers males have relatively smaller testes but higher sperm density and sperm number relative to their testes size (Liljedal and Folstad 2003). Morphological differences can be induced in the same time a dominance hierarchy is established, in just a few days (Liljedal and Folstad 2003), so these morphs are at the extreme of the ephemeral-permanent spectrum.

Måsvær et al. (2004) conducted a correlational study that did not specifically address differences among morphs. Prevalence and intensity of infection of several parasites were combined via Principal Component Analysis. The first 2 principal components correlated with sperm mass and density. However, the PC loadings are not shown, so it is not possible to know whether the relation between parasites and sperm is positive or negative. Nevertheless, no relation was found between immune measures (granulocytes, lymphocytes and spleen mass) and testes size, sperm number or secondary sexual traits.

Liljedal & Folstad (2003) placed males into pairs and a social hierarchy was established in just 4 days. Subordinates were significantly more stressed, but there were no differences in immune function, as measured by circulating granulocyte and lymphocyte counts. Hence, neither stress nor dominance status affected immune condition. It might be illuminating to repeat or expand this work on Arctic char, paying particular attention to morph differences and using experimental infections and more sophisticated immunological techniques.

Immunity and alternative reproductive tactics have been studied in two species of blennies: **The Azorean rock-pool blenny** *(Parablennius parvicornis)*, and the **peacock blenny** *(Salaria pavo).* Both species have sequential polymorphism associated with changes in reproductive behaviour (Santos 1985a; Oliveira et al. 2001). Small young males without secondary sexual characters (M-morph) reproduce as sneakers, and older and larger males develop secondary sexual traits (M+ morph) and follow the so-called bourgeois tactic, either as nest holders or non-territorial floaters (Santos and Almada 1988; Ruchon et al. 1995). The switch is not reversible.

The **Azorean rock-pool blenny** (total length 8-18 cm) inhabits shallow waters on the Atlantic coast of North-East Africa and its adjacent Islands. The best studied population inhabits intertidal pools on Faial, in the Azores. The pools contain crevices that give the fish shelter from most predators. In the spring, large M+ males compete aggressively for these natural cavities (Santos 1985b). Males that manage to defend and clean out a cavity become darker and start courting females (Santos and Barreiros 1993). Females enter the pools at high tide and visit several nest-holding males to lay their eggs. The smaller M-males sneak into the nest when nest-holder males (M+) are inattentive. In contrast, larger M-males settle close to a nest as “satellites”, and provide a service to nest holders (M+) by keeping sneakers (smaller M-males) away. In return, these satellites also benefit, as they manage to fertilize eggs via sneaking when nest-holder males (M+) habituate to their presence. About 30% of Azorean rock-pool blenny males are M- (Fig. 3 of Santos 1995) and most males switch to the M+ morph when they are 2 or 3 years of age (Oliveira et al. 2005).

The **peacock blenny** (total length 5 – 15 cm) breeds in crevices and holes in hard substrate of intertidal areas in the Mediterranean and adjacent Atlantic coast. The best studied population is on Culatra Island in Portugal, where nest-holding males have populated holes in stones and debris that are used to border clam cultures belonging to local fisherman. Nest-holding males (M+) have a conspicuous orange crest and have anal glands. The anal gland produces mate-attracting pheromones and antibacterial compounds that promote egg survival (Pizzolon et al. 2010). M-males are female mimics in appearance and behaviour, and both females and parasitic males (M-) are smaller than M+ males (Gonçalves et al. 2005). The age and size at which M-switch to M+ depends on the availability of suitable nest sites, and when nest sites are abundant, M-males are rarer (Saraiva et al. 2010). At the peak of the reproductive period, 11to 43% (mean 25%) of males are M-(Fagundes, pers.obs.). Females visit several nests and, in an example of mate choice copying, they prefer to lay their eggs in nests that already have large broods (Ros and Oliveira 2009).

Ros *et al.* (2006a) tested the ICHH in rock-pool blennies correlationally by measuring circulating androgens, lymphocyte ratios and primary antibody responses. Both 11-ketotestosterone and testosterone were higher in M+ males than in M-males and lymphocyte numbers and humoral immunity were lower in M+ males than in M-males. However, within each morph, there was no relation between androgens and immunocompetence. Hence, the first result supports the ICHH but the second does not. They suggested that either the hypothesis was incorrect, or that changes in androgens are not necessarily reflected immediately by changes in immunocompetence. Furthermore, they found that in M+ males androgens decreased after an immune challenge (Ros et al. 2006a), which provides some support for the aforementioned “reverse” ICHH.

On the other hand, Ros *et al.* (2006b) conducted an experimental study in which androgens were manipulated and provided partial support for the ICHH. In this study, 11-ketotestosterone, but not testosterone, induced the development of secondary sexual characters in M-males. Furthermore, M-males treated with 11-ketotestosterone swam less than controls. Finally, lymphocytes numbers decreased in testosterone treated males relative to controls, but not in the 11-ketotestosterone group. Hence, the ICHH seems to be too general for the peacock blenny; the androgen that affects morphology and behaviour differs from the androgen that affects immunity. These 2 androgens, 11-ketotestosterone and testosterone, are separated by only one metabolic step. There are no evolutionary or ecological hypotheses of why they have such different effects, or, for that matter, those particular effects. It is unknown whether these differences hold for other species, but this problem might prove to be a fruitful area of research for both ecologists and physiologists.

An energetic constraint explanation is also possible. During the breeding season, nest-holders (M+) spend a lot of resources and time developing secondary sexual traits, defending territories courting females, and cleaning and aerating the nest area, all of which limit their foraging opportunities (Santos 1985a; Almada et al. 1994; Gonçalves and Almada 1997; Ros et al. 2004). In contrast, M-males do not incur these costs and hence, their condition does not decrease as it does in M+ males (Gonçalves and Almada 1997). Hence, differences in androgens between M- and M+ males might not directly affect immunity, but rather regulate the allocation of energetic resources between immunity and reproductive behaviour. As M-males have more future reproductive potential than the older nesting males, young should primarily invest in survival and gradually (as M-) shift that investment to reproduction.

Other explanations are not necessarily mutually exclusive. First, nest-holders (M+) aggressively defend the nest against parasitic males (M-), and while doing so they inflict typical scratch-like injuries with their large canine-like teeth. In Azorean rock-pool blennies, M-males have many more such injuries than nest-holder M+ males. As a consequence M-males are more exposed to potential infections with pathogenic organisms, which might explain why they are more immunocompetent (Ros et al. 2006a). Second, one might expect older males to have been exposed to most of the common pathogens in their environment and to have acquired immunity. As a result older (M+) males might decrease production of naïve leukocytes which would be necessary for acquiring immunity to new pathogens. Therefore, the fact that if M+ males had lower lymphocyte blood cell counts and lower antibody responsiveness than M-males does not necessarily mean that they are more susceptible to disease (Ros and Oliveira 2009). Under this scenario, the immune system improves with age, but using some measures of immunity, the exact opposite is true. Longterm studies in which related individuals are raised in different pathogen and social environments are needed to separate these hypotheses.

**The Corkwing wrasse** (*Symphodus melops*) is an intertidal species occurring in the western Mediterranean and Adriatic seas, and in the Eastern Atlantic, from Morocco and the Azores in the South, to Norway in the North. They mostly live in waters less than 5 m in depth. Adults are 15-25 cm long. Females and juveniles are greenish brown and territorial males are iridescent blue-green. Territorial males build and defend nests where several females might deposit their eggs. In addition, between 2-20% of males are female mimics, which do not build or defend nests. Female mimics resemble females to the point of having a urogenital papilla similar to that of females. Corkwing wrasses do not change sex, nor do they change male morph. Female mimics of all ages and sizes occur, so it is not a sequential, age- or size-dependent polymorphism. It is unknown whether the male morphs are strictly genetically determined, or environmentally determined early in the males’ lives. Female mimics are smaller than regular males but have relatively larger gonads, more motile and longer-living sperm than territorial males (Potts 1974; Dipper and Pullin 1979; Dipper 1981; Quignard and Pras 1986; Costello 1991; Darwall et al. 1992; Uglem et al. 2000).

Uglem *et al.* (2001) tested the sperm protection hypothesis and found that territorial males have smaller gonads and larger spleens than female mimics, but the density of lymphocytes and granulocytes did not differ between the two male morphs. So, the cell counts did not support the hypothesis, and the differences in spleen size were the exact opposite of what is predicted by the sperm protection hypothesis (Uglem et al. 2001). The larger gonads of sneaker morphs are as expected based on sperm competition. Subsequent work confirmed that territorial males have larger gonads and more motile sperm than female mimics, but sex steroids affect only sperm quantity, and not sperm quality (Uglem et al. 2002). Hence, contrary to the sperm protection hypothesis and the ICHH, in this system was no detectable relation between sperm quality, immunity, and hormones.

### 4.2 Birds

Plumage polymorphisms in birds are fairly common (Galeotti et al. 2003), and are maintained by a variety of processes (Lank 2002). However, ARTS, as defined in this review, that is, relatively longterm and literally “alternative”, occur in only seven species (Krüger 2008). Although much immunoecological work has been conducted with birds, so far, it has only been conducted on two species with ARTS.

The **ruff** is a lekking shorebird that breeds in marshes and grasslands in northern Eurasia and winters mostly in Africa. There are three morphological and behavioural types of males: independents, satellites and faeders. Independents and satellites have elaborate plumage and courtship behaviour (Hogan-Warburg 1966; van Rhijn 1991), and faeders are female mimics (Jukema and Piersma 2006). About 1% of males are faeders (Verkuil et al. 2008), 15% satellites, and the rest independents. Ruffs range in mass from 70 g to 220 g, with extensive overlap between the morphs and sexes. Independents are the largest and females the smallest; faeders are at the upper end of the female distribution and the lower end of the male distribution (Bachman and Widemo 1999; Lank et al. 2013). Females carrying the faeder gene are the smallest (Lank et al. 2013). The faeder-like morph arose from the ancestral independent about 3.8 mya, and the satellite 0.5 mya (Kupper et al. 2016; Lamichhaney et al. 2016). Females carry the independent/satellite alleles but express the respective phenotypes only when experimentally exposed to testosterone (Lank et al. 1999).

Faeders migrate to the breeding grounds later that most other males, but earlier than most females (Karlionova et al. 2007). During the breeding season, independents establish lek mating courts defend them against each other but try to attract into their courts satellites and females (Höglund and Lundberg 1989; Hill 1991; van Rhijn 1991). Satellites do not fight but are pecked and chased by independents when they fail to be submissive and when they attempt to mount a visiting female, but they are tolerated because co-occupied courts are more likely to attract females (Höglund and Lundberg 1989; Hill 1991; van Rhijn 1991). Faeders look like and behave like females, spending time among females and visiting leks unchallenged. Nevertheless, faeders readily copulate when females solicit copulations from independents, satellites or perhaps even faeders.

Lozano and Lank (2004) tested the ICHH and the energetic constraints hypothesis in a captive population. They examined whether early in the breeding season immunity was positively correlated size and the degree of development testosterone-induced sexual traits. This work was conducted before the discovery of faeders. Cell-mediated immunity (CMI) was estimated using a delayed hypersensitivity test and humoral immunity using a SRBC agglutination test. However, other than the fact that CMI was weakly correlated with ruff length, neither measure of immunity was related to any other sexually selected traits, size of mass, in either satellites or independents. Lozano and Lank (2004) also tested the energetic trade-off hypothesis, which predicts that satellites should have stronger immune responses than independents. Although humoral immunity did not differ between the two male morphs, independents had **higher** CMI responses than satellites. If anything, these results support the “risk of injury” hypothesis. In contrast, during the non-breeding season the immune responses of both morphs were stronger, but despite a larger sample size (51 vs. only 21 males in the breeding season), there were no differences in CMI between the morphs (Lozano and Lank 2003). Hence, the decrease in immune function during the breeding season compared to the non-breeding season supports the energetic constraints hypothesis, but differences in immunity between independents and satellites **during** the breeding season are best explained by “risk of injury” hypothesis.

These two alternatives were subsequently tested by Lozano *et al.* (2013). This latter study included the recently discovered faeder, the female mimicking male morph. Based on their behaviour and life history, the 3 male morphs and females can be placed on an ordinal scale with independents at one end and females at the other. A haemolysis-haemagglutination assay (Matson et al. 2005) on samples taken before and after an injection with liposaccharide was used to measure innate and adaptive humoral immunity, respectively. Furthermore, CMI was once again estimated via a delayed hypersensitivity test. No significant differences were evident between the four groups, but all three measures of immunity decreased along this axis from independents to females. These results support the risk-of-injury hypothesis over the energetic constrains hypothesis. Hence, the immune responses of the 4 ruff genders reflected their life history and behaviour, with faeders located in the immunological continuum between females and the other male morphs. So far in this system, support is lacking for hypotheses predicting a direct hormonal regulation of immune responses, as per the ICHH and its variants.

Additionally, in ruffs CMI decreases significantly with age, in males (Lozano and Lank 2004), and in both sexes (Lozano and Lank 2003). Nebel *et al.* (2013) partially confirmed these results, although only for females, and with only one of several immune measures. Although the decrease was not related to morph differences, it is consistent with the basic life-history premise that investment on self-maintenance should decrease as residual reproductive success wanes. Finally, again, although morph differences were not detected, Lozano & Lank (2003) showed that younger birds have weaker immune responses, presumably because of the way a neonate’s immune system develops, by learning which antigens are harmful, and/or because of energetic constraints as a neonate’s resources are channelled primarily towards growth.

The **white-throated sparrow** *(Zonotrichia albicolis)* is a common songbird that breeds in the northeast of the USA and Canada east of the Rockies, and winters mostly in southern and eastern USA. Both sexes have two morphs, defined by the colour of the medial crown stripe (a stripe on their heads): tan striped (TS) and white-striped (WS) (Lowther 1961). The morphs are caused by a chromosomal inversion, and hence, are fixed (Thorneycroft 1966; Throneycroft 1975). Birds mate with the opposite morph, which maintains the population frequencies of the two morphs roughly equal (Houtman and Falls 1994).

At the wintering grounds, WS birds, particularly males, are more neophobic than TS birds (Barcelo-Serra et al. 2020). In contrast, WS birds are more aggressive than TS birds on the breeding grounds (Horton et al. 2012). Additionally, both male and female WS birds sing more than their TS counterparts (Collins and Houtman 1999). Furthermore, compared to TS males, WS males respond more aggressively to territorial intrusions (Collins and Houtman 1999), are less parental (Knapton and Falls 1983), and are more likely to engage in extra-pair copulations (Knapton and Falls 1983; Tuttle 2003). Additionally, female WS are more likely to solicit EPCs and dump their eggs in neighbouring nests (Tuttle 2003). Whereas the genetic and endocrine mechanisms causing and associated with the polymorphism have been extensively studied (e.g., Spinney et al. 2006; Maney 2008; Huynh et al. 2011; Hedrick et al. 2018), so far there is only one study addressing immunological differences.

Boyd et al. (2018) looked for sex and morph differences in immunity under the general guise of “extended immunocompetence handicap models”, predicting lower resistance in the WS than the TS morph to haemosporidian parasites. Forty-two birds, 23 males (15 WS and 9 TS), and 19 females (7 WS and 12 TS) were captured after the breeding season, during the fall migration. Twelve of these were already infected, but there were not differences between the two morphs. They were maintained in captivity for 4 months in a short day photoperiod (10 h Light, 14 h Dark). On the fifth month, they were switched to a long-day photoperiod (16 L: 8D) and inoculated or sham-inoculated with *Plasmodium* parasites obtained from two donors. Of the 43 birds, 12 were sham-inoculated and 30 inoculated. Of the 30 inoculated birds, two thirds (21) were completely resistant. Of the 9 infected birds (4 TS, 5 WS), the two highest parasite loads belonged to TS individuals. Androgens were higher in males than in females, as it would be expected, but they did not differ between the morphs. Therefore, no sex differences were detected, and if anything, parasite loads ended up being higher in TS than WS birds, contrary to the main prediction.

The study clearly suffered from a small sample size, but the most puzzling aspect is why the birds were captured at the end of the breeding season and kept in captivity for 4 months before conducting the study. The most notable result, that two thirds of all birds were completely resistant to *Plasmodium* infections, was not discussed. Boyd et al (2018) concluded that their results did not support the mechanistic androgen-dependent ICHH, but did not consider ecological hypotheses. Given that WS were more aggressive and likely suffered greater exposure to parasites than TS birds, the “risk of injury” hypothesis could have been considered.

### 4.3 Reptiles

A variety of ARTS exist among reptiles. These include r- and k-life histories, manipulation of offspring sex, territoriality vs. non-territoriality, parthenogenesis vs. sexual reproduction, and behavioural, morphological and chemical female mimicry (Crews 1983; Calsbeek and Sinervo 2008). Unlike fish, reptiles lack the ability to change sex during adulthood, so the range of reproductive tactics and strategies does not include all of the ones present in fish, but they do include all of the ones present in birds. With reptiles, one must be careful to distinguish between merely polymorphic species and species with ARTS, as defined above.

Several studies have compared immune function of different morphs in polymorphic species. For instance, work was been conducted on the brown anole *(Anolis sagrei)* (Calsbeek et al. 2008), the wall lizard *(Podarcis muralis)* (Sacchi et al. 2007; Galeotti et al. 2010), and Iberian wall lizard *(Podarcis hispanicus)* (Ortega et al. 2015). In these studies, the general rationale seems to be that differences in immune responsiveness, presumably reflecting differences in parasite exposure or susceptibility to disease, contribute to the maintenance of the polymorphism. However, the reasons are not explicitly stated, and the predictions are nebulous. Even when significant differences are found, it is unclear why one morph’s immune response would be stronger than the other morph’s. In contrast, when using species with ARTS, or merely when considering the morphs’ ecology, it is possible to articulate specific directional predictions and design more purposeful tests.

The **Dalmatian wall lizard** *(Podarcis melisellensis)* is endemic to the Adriatic countries. Adults have an SVL of at least 5 cm; they lay 3 clutches per year with 3-9 eggs per clutch, and have a clearly evident trade-off between egg size and number. Males have a SVL of about 6 cm, their ventral surface can be orange, yellow, or white. They do not change between successive breeding seasons, so it is most likely that the morphs are permanent and established early in life. Although their sizes overlap considerably, orange males are generally larger and more aggressive than either white or yellow males (Huyghe et al. 2007).

Huyghe *et al.* (2009) compared testosterone, corticosterone, morphology, biting strength, and immunity among the three morphs. The study framed around the immunocompetence handicap hypotheses, and it was predicted that aggressive orange males would have lower immunity and hence higher parasite susceptibility than the other morphs. Immunity was measured using the aforementioned delayed hypersensitivity test, and by counting the numbers of mites and ticks. Nevertheless, no significant differences were found among morphs in neither testosterone nor either measure of immunity.

Subsequently, Huyghe *et al.* (2010) examined seasonal differences in cell-mediated immunity in males, again using PHA test for cell mediate immunity, ectoparasites, and blood parasites. Again, the study framed around the immunocompetence handicap hypotheses, perhaps mixed with the energetic trade-off hypothesis, and it was predicted that, as the season progressed, aggressive orange males would have lower immunity and hence higher parasite susceptibility than the other morphs. Ectoparasite prevalence was higher at the end of the breeding season than at the start for all morphs, but there were no differences among the morphs. The prevalence of blood parasites increased across the season for white and yellow morphs, but decreased for the orange morph. Unfortunately, data for parasite intensity are not shown; instead, the exact same figure, the one for prevalence, is presented twice (Figs. 1 and 2 in Huyghe et al. 2010). PHA responses followed the same pattern as prevalence; compared to the start of the breeding season, at the end of the season PHA responses had increased in white and yellow males, but it remained unchanged in orange males. Hence, no support was found for the ICHH or the energetic trade-off hypotheses, but perhaps some support for the exposure risk hypothesis.

The **side-blotched lizard** *(Uta stansburiana)* is a small (50-60 mm SVL, 7-10 g), highly polymorphic lizard common in the south-west of North America (Tinkle 1967). It sometimes reaches high densities, of up to 2,600 per hectare (Svensson et al. 2001). They mature at one year of age, and usually live for only one breeding season. The species is separated into several populations or subspecies, usually with different types of polymorphisms (Upton and Murphy 1997).

In the population studied by Sinervo and colleagues, used for all the studies below, morphs are determined by 3 alleles (o, b, and y). Males occur in three morphs: orange, yellow, and blue. In males, the “o” allele is dominant, and the “b” allele is recessive to the “y” allele (oo, ob, and oy are orange, yb and yy are yellow, and bb are blue). Orange males are aggressive and highly territorial, blue males have smaller territories and are not as aggressive and yellow males are sneakers. Male morphs are maintained via frequency dependent section in a “rock, paper, scissors” system (Sinervo and Lively 1996), whereby orange males can take over territories of blue males, but are themselves vulnerable to cuckoldry by yellow males (Sinervo and Lively 1996). Females occur in two morphs, orange and yellow. In females, the “o” allele is also dominant (oo, ob, and oy are orange, yb, yy, and bb are yellow) (Sinervo and Zamudio 2001). Orange females produce large clutches of small eggs and are favoured at low densities, and yellow females produce small clutches of large eggs and are favoured at high densities. Their relative abundances oscillate in 2-year cycles, driven by negative frequency dependent selection.

Using antibody responsiveness towards a tetanus toxoid antigen, Svensson *et al.* (2001) found that population density was negatively related with immune responsiveness of both morphs but more so in orange females. Furthermore, subsequent survival was positively related to the strength of prior antibody responsiveness in yellow females, but negatively related in orange ones (Svensson et al. 2001). Hence, compared to yellow females, orange females, which are adapted to fast reproduction, invest less in immune function, and are less capable of activating their immune system without suffering negative consequences. These trade-offs are as expected based on life history theory, but seldom are they documented within one sex.

Svensson *et al.* (2009) extended these findings to include males, examining the population genetics behind colour and humoral immunity. They found orange colour has highly heritable (0.46), and antibody responsiveness to diphtheria-tetanus antigens even more so (0.61 for sons and 0.82 for females). However, the correlations between orange colour and antibody responsiveness were in opposite directions for males and females, producing a significant intersexual conflict (Svensson et al. 2009). This means that, assuming there is no post-copulatory mate choice, females cannot simply select males based on good immunity, but rather must choose between producing immunocompetent sons or immunocompetent daughters. The implications are yet to be examined.

### 4.4 Insects

As it might be expected for such a speciose and diverse taxon with distinct life stages, a wide variety of ARTS occur among insects (Brockmann 2001, 2008). Brockmann (2008) divided reproductive behaviour in insects into four sequential events: (1) locating mates (2) obtaining access to mates, usually females, (3) copulating, and (4) guarding the female after copulation. Except for the very act of copulation, ARTS occur at all stages. Although the literature on ARTS in insects is extensive, and their immune systems relatively simple compared to that of vertebrates, it is peculiar that few researchers have examined the immunoecology of insects with ARTS.

The beetle *Onthophagus taurus* is a member of the **scarabaeid dung beetle** group, a taxon in which only males have horns. Diet during the larval stage determines whether individuals reach a certain size threshold at a given time in their development, at which point two growth trajectories are followed. Males above the size threshold develop unusually large horns and males below the threshold develop only rudimentary horns. This dimorphism is associated with behavioural dimorphisms. Horned males compete directly with each other in their tunnels for access to females, and guard them after copulation. Hornless males are sneakers and dig across tunnels, enter the tunnels beneath the guarding male and, and undetected by the guarding male, mate with females (Emlen 1994; Emlen 1997; Moczek and Emlen 1999). Both sexes excavate branched tunnels under dung patties, where females lay their eggs and leave some dung for their offspring to eat when they hatch. However, horned males provide more paternal care than hornless males, but only when there are no other males present (Moczek 1999).

Cotter *et al.* (2008) examined immunity in male larvae, after the eventual morph had been determined but before the actual development of horns. At that point in their development, their diet has been slightly different, their size differs, but energetic requirements do not differ much yet. Unfortunately, trade-offs are mentioned but no hypothesis is explicitly addressed, so the expectations are unclear. Nevertheless, using phenoloxidase (PO) activity as a measure of innate immunity, they showed that, controlling for condition, male larvae that will eventually be horned have higher innate immunity than larvae of future hornless males. Perhaps these differences are a prophylactic investment on immune function and reflect future differences in pathogen exposure. Female PO activity was between that of the two male morphs, which would be potentially interesting if there were more explicit hypotheses predicting such a pattern. Because horn growth is induced by juvenile hormone (Emlen and Nijhout 1999), Cotter *et al.* (2008) suggest that juvenile hormone also causes the differences in immunity. This juvenile hormone hypothesis does not depend on energetic trade-offs (proximate or ultimate), but it does provide a mechanism regulating immunity in insects, akin to the suggested androgen mechanisms for vertebrates. This idea is yet to be tested.

**Odonates** present an interesting contrast in that females, not males, are the ones with alternative morphs. In 134 out of 195 species, there is one or more females morphs (called gynomorphs or gynochromes) and a male mimicking morph (called andromorphs or androchromes) (Fincke et al. 2005). Andromorphs mimic both male morphology and behaviour. Compared to gynomorphs, andromorphs are brightly coloured and have narrower abdomens, and they are more aggressive and more likely to roam about in open spaces (Robertson 1985). The polymorphism is based on a female-restricted autosomal system with 2 or 3 alleles (Johnson 1964; Johnson 1966; Andrés and Cordero 1999). The proportion of andromorphs is highly variable, but generally, polymorphisms are more common in populations with relatively more males than females (van Gussum et al. 2007; Iserbyt et al. 2009). For example, in *Ischnura elegans*, depending on the population, andromorphs constitute 14% to 94% of all females (Sánchez-Guillén et al. 2013a). In *Nehalennia irene* (Hagen) andromorph frequencies range from 0 to > 90% across Canada (van Gussum et al. 2007).

Andromorphs are subjected to less sexual harassment than gynomorphs. For instance, in *Ischnura senegalensis*, males harass gynomorphs more than they do andromorphs; compared to andromorphs, gynomorphs do not defecate as much, which presumably reflects their food intake, and most importantly, they produce 35% fewer eggs (Takahashi and Watanabe 2010). In *Ischnura elegans*, andromorphs mate less often than gynomorphs, and males are more attracted to gynomorphs (Cordero et al. 1998). As usual, the picture is not so simple. Another study of *Ischnura elegans* showed that naïve males prefer andromorph over gynomorph females, but experienced males have no such preference (Sánchez-Guillén et al. 2013a).

Although the consensus is that alternative female morphs are maintained, and perhaps have evolved because they help females decrease male harassment, that argument just reverses the question. Unless there is a cost to being an andromorph, andromorphs would eventually become fixed in a population. Sánchez-Guillén *et al.* (2013b) suggested that andromorphs are more conspicuous and more exposed to parasites than gynomorphs, so increased parasitism might counterbalance the benefits of reduced male harassment: the “exposure risk” hypothesis. They tested the idea by quantifying the prevalence of intestinal gregarines (Apicomplexa: Eugregarinidae) and aquatic water mites *(Arrenurus sp.)* in several species of damselflies. Gregarine prevalence did not differ between the morphs, but, as predicted, mite prevalence was higher in andromorphs than in gynomorphs.

Joop *et al.* (2006) conducted a similar study using the azure damselfly *(Coenagrion puella).* Once again, the general rationale is that differences in immune responsiveness contribute to the maintenance of the polymorphism, but the reasons are not explicitly stated. A link to melanin is mentioned, curiously, more prominently in the abstract than in the body of the paper, but the link is not explained. The andromorph is blue, the gynomorph is green, and as one might expect, the intermediate is blue-green. These colours are more likely to be structural, not melanin dependent, and nothing in the paper indicates otherwise. Nevertheless, no differences were found among the morphs in the prevalence of water mites or gregarines. Furthermore, following an experimental fungal infection, there were no differences in resistance among female morphs.

## 5 CONCLUSIONS

Males and females are dissimilar in so many ways that it would not be surprising to find one more difference. The core benefit of testing hypotheses using species with ARTS is that these species have additional genders beyond the usual two, so it is possible to disentangle the effects or sex *per se* from the effects of various morphological and behavioural adaptations that are usually associated with sex. Hypotheses predicting differences between two sexes could actually be more thoroughly tested by taking advantage of the additional morph(s) because any carefully articulated hypothesis would yield falsifiable, *a priori*, directional predictions. To be able to separate such effects, it is of paramount importance that the hypotheses are carefully delineated, and the predictions clearly articulated. This would be true for any hypotheses, not just those related to immunoecology. Unfortunately, this problem is evident in several immunoecological studies using ARTS, whereby the hypotheses and predictions are not clearly articulated. Instead, studies often just test for differences among morphs, hence squandering the main advantage of working with species with ARTS.

The paucity of work on insects is surprising. Let us be clear. Plenty of work exists on insect polymorphisms, and some of it on immunoecology; however, we were not able to find many immunoecological studies in species with alternative life histories. There are myriad benefits to working with insects. Generally, the sheer number of species of insects, and arthropods in general, means that researchers could find the best possible species with which to test a particular hypothesis. Second, given their small size, obtaining a reasonable sample size becomes less of a problem. Also, it goes without saying that it is easier to get permits to work with insects. There are also other benefits to working with insects more specifically relevant to immunoecology in species with ARTS. First, many types of polymorphisms, both morphological and/or behavioural, have already been described in insects (Brockmann 2008), and many of these are technically “alternative”. Second, compared to vertebrates, their immune systems are relatively simple, so it might be easier to focus on the ecology and avoid being bogged down by the mechanisms of immunity. For instance, their immune system does not have immunological memory (Söderhäll 2010); therefore, should experimental design require it, repeatedly assessing the immune systems of individuals would not compromise the study’s validity. Hence, researchers who are truly interested in trade-offs between maintenance and reproduction in species with alternative life histories, and who are not required to work on vertebrates, would do well by choosing to work with insects.

For 30 years now, the ICHH, has been a starting point for many studies, but maybe it has had too much influence. In several cases, it is considered as a generic starting point and explicit predictions are not clearly articulated. Meanwhile, other hypotheses have been relatively ignored. In several cases, after failing to find support for the ICHH, authors fall back on these hypotheses *a posteriori.* Instead of being relegated to the end of the paper as caveats and unexplored possibilities, perhaps is it time for these physiological, ecological, life history hypotheses to take centre stage, or at least be considered *a priori*, even in conjunction with the ICHH.

Even if the ICHH it is still considered a convenient starting point, it is time to examine the subtleties within it. For instance, by examining several indices of immunity and several androgen hormones, studies can increase the chances of finding some support for the hypothesis. However, different androgens often have different effects on different aspects of the immune system, but there are never any evolutionary, ecological, or phylogenetic explanations of why some androgens, but not others, should affect immunity in that particular way. It is at this point that the unification of the ecological and the physiological approaches disintegrates and the focus is shifted to physiological mechanisms. In immunoecology, as in any branch of physioecology, we must remain keenly aware that the proximate and ultimate approaches pull us in opposite directions. It seems that evolutionary ecologists are easily seduced by the sophisticated instrumentation and technical complexity of the reductionist approach, in the process forgetting the evolutionary history and ecological intricacies of their beloved organisms.

In evolutionary ecology, hypotheses must not merely deal with physiology and mechanisms but also address the far more relevant question of how ecological and evolutionary forces have produced these particular mechanisms. One possible fruitful way to merge the two approaches might be to start with how ecology and life-history not only affect immunity, but more specifically, when and how they would activate and/or suppress immune function. Many papers focus on immunosuppression or immunotolerance and give adaptive explanations for this in terms of tradeoffs or in terms of energetic constraints. However, all these could be divided into two general situations that might determine when to: 1) activate or stimulate immunity, for instance, to fight a new pathogen, and 2) suppress or subdue it, for instance, to temporarily tolerate a parasite. A suitable regulation mechanism is one that is flexible enough so it can respond to ecological opportunities, both during an individual’s lifetime and at evolutionary time scales (Hau 2007). Thus the trade-off between immunocompetence and reproduction will change with age and with the particular morph. Similar conceptual frameworks merging the ultimate and proximate approaches might be necessary for continued development of immunoecology, and in particular when dealing with species with ARTS.

Interactions between hosts and parasites are yet to be fully examined in the evolution of ARTS. Parasites, and hence immunity, might also play a role in the evolution of ARTS (Todd 2007). Alternative morphs may arise because of differential parasite resistance strategies, especially when parasite resistance is modulated by a hormone that is also involved in reproductive investment (Zera and Harshman 2001). Empirical work on which the ICHH and its derivatives are based documents the existence of such trade-offs. Given these trade-offs, dominant males may be more susceptible to the parasites, which would lower their ability to defend their reproductive resources and sexual partners.

Furthermore, by definition, parasites have the potential to decrease their hosts’ fitness, but many parasites co-evolve with their hosts and reduce their virulence but increase their reproduction. Consequently, older hosts might be more infected and more chronically infected than younger hosts. The burden of parasitism might lower the condition of dominant males and facilitate the evolution of ARTS.

Finally, several recent studies have specifically addressed genomics in species with ARTS (Pointer et al. 2013; Schunter et al. 2014; Stuglik et al. 2014). Species with ARTS are clearly great systems in which to examine sex and morph-specific gene expression. Using these rapidly advancing techniques, it is now possible to identify large number of genes involved in the development of morphs and the regulation of immunity, and conduct truly integrative studies across all levels of complexity. However, no technical advance is without its conceptual detractors. Because of polygenic effects on phenotypes, the reciprocating modifying effects of organisms and their environments, and the fact that several evolutionary and ecological selection regimes can produce a given genome, it has been argued that further genomic details will distract us from our goals and detract from our understanding (Travisano and Shaw 2013). Others acknowledge the pitfalls, but advocate cautious optimism (Zuk and Balenger 2014). In the case of ARTS, knowing the genetic basis of morph development and immune regulation within each morph might provide unexpected insights, but without ecological and evolutionary hypotheses driving the effort, we will not really know how immunity interacts with the evolution and maintenance of these morphs. Decoding entire genomes and disentangling physiological mechanisms can be informative, but there is no grandeur in the strict reductionist view of life.

## ACKNOWLEDGEMENTS

G.A.L. thanks the University of Tartu library for allowing him free access to their online collections and L. Lozano (née Prigo) for getting books from the said library under her name. This is paper contribution number 2114 of the ECEE. We appreciated the comments Corinna von Kürthy made on a previous version.

## Footnotes

1 After Mithridates VI (135 – 63 B.C.), King of Pontus (113 – 63 B.C.), who, legend has it, purposely exposed himself to poisons in an attempt to become immune to them (Mayor 2009).

